# Automated Visualization of Rule-based Models

**DOI:** 10.1101/074138

**Authors:** John A. P. Sekar, Jose-Juan Tapia, James R. Faeder

**Affiliations:** University of Pittsburgh School of Medicine Department of Computational & Systems Biology

## Abstract

Rule-based modeling frameworks provide a specification format in which kinetic interactions are modeled as “reaction rules”. These rules are specified on phosphorylation motifs, domains, binding sites and other sub-molecular structures, and have proved useful for modeling signal transduction. Visual representations are necessary to understand individual rules as well as analyze interactions of hundreds of rules, which motivates the need for automated diagramming tools for rule-based models. Here, we present a theoretical framework that unifies the layers of information in a rule-based model and enables automated visualization of (i) the mechanism encoded in a rule, (ii) the regulatory interaction of two or more rules, and (iii) the emergent network architecture of a large rule set. Specifically, we present a compact rule visualization that conveys the action of a rule explicitly (unlike conventional visualizations), a regulatory graph visualization that conveys regulatory interactions between rules, and a set of graph compression methods that synthesize informative pathway diagrams from complex regulatory graphs. These methods enable inference of network motifs (such as feedback and feed-forward loops), automated generation of signal flow diagrams for hundreds of rules, and tunable network compression using heuristics and graph analysis, all of which are advances over the state of the art for rule-based models. These methods also produce more readable diagrams than currently available tools as we show with an empirical comparison across 27 published rule-based models of various sizes. We provide an implementation in the open source and freely available BioNetGen framework, but the underlying methods are applicable to all current rule-based models in BioNetGen, Kappa and Simmune frameworks. We expect that these tools will promote communication and analysis of rule-based models and their eventual integration into whole cell models.

**Author Summary:** Signaling in living cells is mediated through a complex network of chemical interactions. Current predictive models of signal pathways have hundreds of reaction rules that specify chemical interactions, and a comprehensive model of a stem cell or cancer cell would be expected to have many more. Visualizations of rules and their interactions are needed to navigate, organize, communicate and analyze large signaling models. In this work, we have developed: (i) a novel visualization for individual rules that compactly conveys what each rule does, (ii) a comprehensive visualization of a set of rules as a network of regulatory interactions called a regulatory graph, and (iii) a set of procedures for compressing the regulatory graph into a pathway diagram that highlights underlying signaling motifs such as feedback and feedforward loops. We show that these visualizations are compact and informative across models of widely varying sizes. The methods developed here not only improve the understandability of current models, but also establish principles for organizing the much larger models of the future.

## Introduction

Biochemical systems are information processing architectures whose building blocks are chemical interactions [1,2]. Predictive models of biochemical signaling are typically in the form of reaction networks [1,2]. In the reaction network specification, a model consists of chemical species (molecules and complexes) and reactions which transform reactant species into product species at specific rates [1,2]. However, in a real biochemical system, information is embedded not just in the identity of each chemical species, but in the interactions of sub-molecular components such as phospho-motifs, domains and binding sites that can be configured in a modular fashion [3]. This has negative implications for the reaction network specification: model hypotheses are hard to recover (Fig 1A) and highly resolved signaling models require disproportionately large reaction networks (Fig 1B). To address these shortcomings, a number of frameworks have emerged that specify and simulate chemical kinetics using “reaction rules”, i.e. reaction classes specified explicitly at the sub-molecular level (Fig 1A). These frameworks are collectively called rule-based modeling and a number of rule-based models have been published in recent times (see [4] for a review and S6 Appendix for a partial list). Model sizes currently range from tens to hundreds of rules, but these numbers are expected to increase as rule-based models are eventually integrated into whole cell models (e.g. [5]) and reaction rules are collectively organized in databases of kinetic interactions [6–9]. Automated visualization tools scalable to large numbers of rules are necessary for rule-based models to be transparent, reusable and comprehensible to a wider audience, but currently available visualization tools are either manual, incomplete or scale poorly with number of rules (see Discussion). In this work, we provide a unified framework for visualizing rule-based models at different levels of resolution that is scalable to large numbers of rules. We address three commonly encountered visualization goals: what chemical interaction does each rule represent, how does information flow between rules, and what signaling architecture does the model represent?

**Fig 1.**
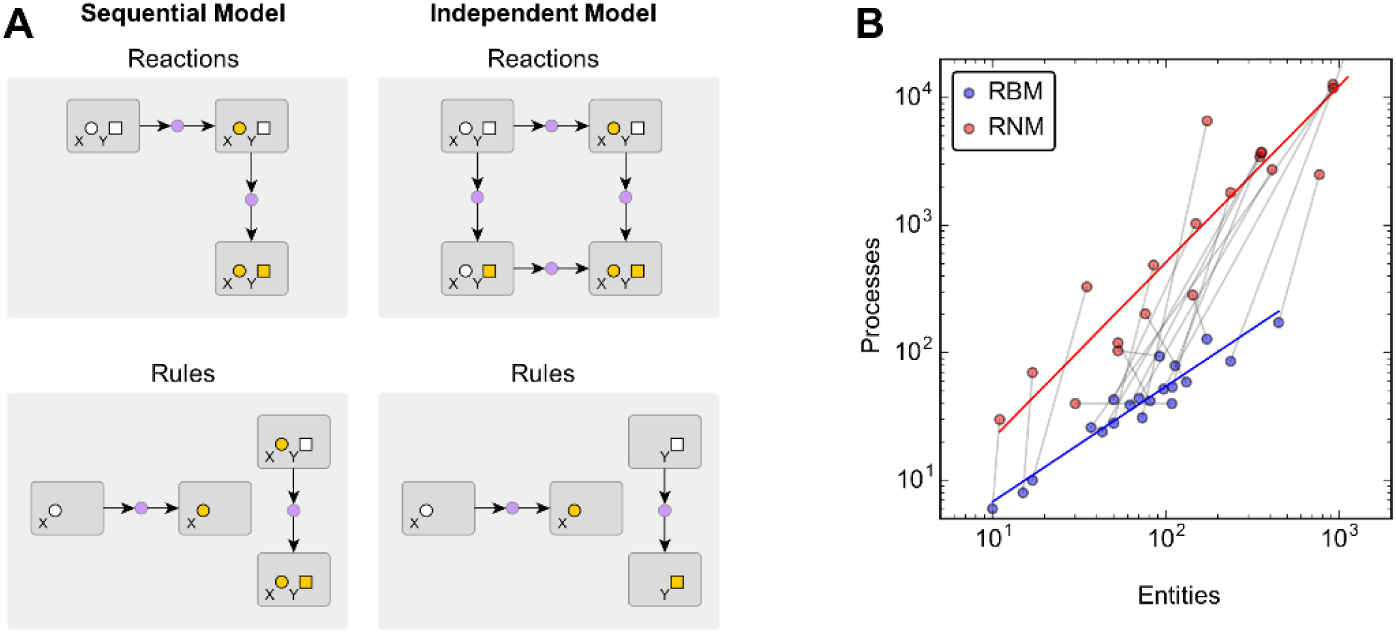
Motivation for rule-based modeling. **(A)** Two models (sequential & independent) encoded using two different specifications (reactions & reaction rules). In the sequential model, site X is activated independently, but site Y is activated only when X is active. In the independent model, both sites are activated independently. When specified as a reaction network, each chemical species and reaction is built and verified manually. Model assumptions manifest as global network properties: the X->Y dependence in the sequential model is verified by checking for the absence of an alternative activation route for Y. When specified as a rule-based model, only partial species and reactions are specified and they are called patterns and reaction rules respectively. Model assumptions are encoded locally by explicitly choosing the sites specified in a rule. In the sequential model with rules, the first rule encodes that X is activated independently of Y since Y is not specified in the rule. The second rule encodes that Y is activated when X is active, since active X is specified on both sides of the rule. In the independent model, both rules encode independent activations, since they do not specify any site other than the modified one. **(B)** Comparison of specification sizes for 20 rule-based models (RBMs) from the literature and their equivalent reaction networks (RNMs). Entities and processes are patterns and rules respectively in RBMs (blue) and species and reactions respectively in RNMs (red). The gray lines link each rule-based model with its corresponding reaction network. The reaction networks are much larger because they grow as the number of possible combinations of sites. However, the number of reaction rules only grows as the number of model assumptions about kinetic dependence, which is a much smaller number.

The building blocks of a rule-based model are the pattern and the reaction rule, just as the building blocks of a reaction network are the chemical species and the reaction. The pattern is a graph that is used to specify sub-molecular structural features (Fig 2A) and specifying a pattern is equivalent to specifying criteria for dividing the state space of chemical species into matching and non-matching subsets [10–12]. A reaction rule is composed by using patterns as reactants and products (Fig 2B) and assigning a rate law. Specifying a reaction rule is equivalent to specifying similar reactions on all combinations of species matched by the reactant patterns and assigning a rate law to each reaction [10–12]. Information flow in a rule-based model is a consequence of features shared between patterns across different rules, for example, when the action of one rule produces a structure that is required by another rule (Fig 2C). By organizing the facts about sites in the model, the reaction rules that operate on them and the overlaps between the rules, one can infer the signal architecture of the model (Fig 2D). Thus, it is useful to organize a rule-based model under four hierarchical layers: structure (patterns and matched species), mechanism (reaction rules and matched reactions), regulation (overlaps between rules) and function (signal architecture). The three visualization goals stated previously correspond to the visualization of the mechanistic, regulatory and functional layers respectively.

**Fig 2.**
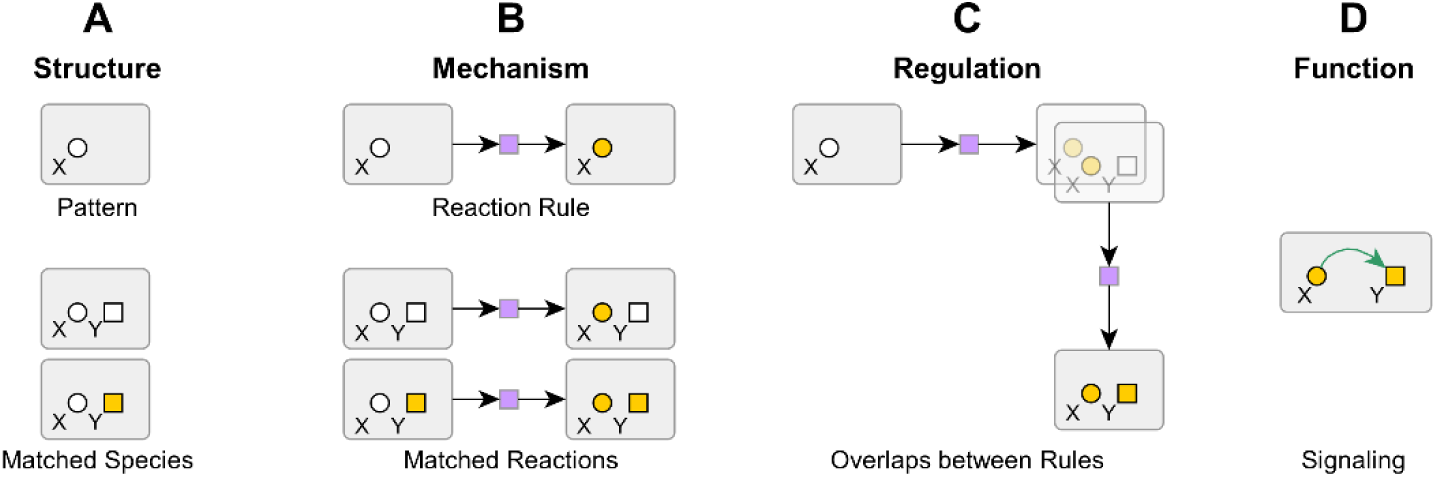
Layers of information in a rule-based model. **(A) Structure.** Each pattern (partially specified species) matches a class of chemical species. **(B) Mechanism.** Each reaction rule (chemical interaction defined on patterns) matches a class of similar reactions. **(C) Regulation.** Overlaps between patterns across rules mediate information transfer (e.g. X is activated in one rule and required in another). **(D) Function.** The facts of the model can be organized in the form of a signaling architecture (e.g. X activates Y).

A number of visualization tools could potentially be applied to rule-based models, but they are fragmented across different fields and have been defined or implemented for rule-based models to different extents. Some of these approaches include conventional rule visualization [13–15], contact map [16], rule influence diagram [17], Kappa story [18], Simmune Network Viewer [19], the Systems Biology Graphical Notation (SBGN) [20], the Molecular Interaction Map (MIM) [21], the Extended Contact Map (ECM) [14] and the *rxncon* regulatory graph [22]. In Fig 3, we apply four of these tools to a previously published model of signaling through FcεRI receptors in mast cells [13], with each tool representative of a particular layer of information. Using these, we can demonstrate the problems with the current state of the art and the lack of an integrated visualization framework. We address the remaining tools in the Discussion section.

**Fig 3.**
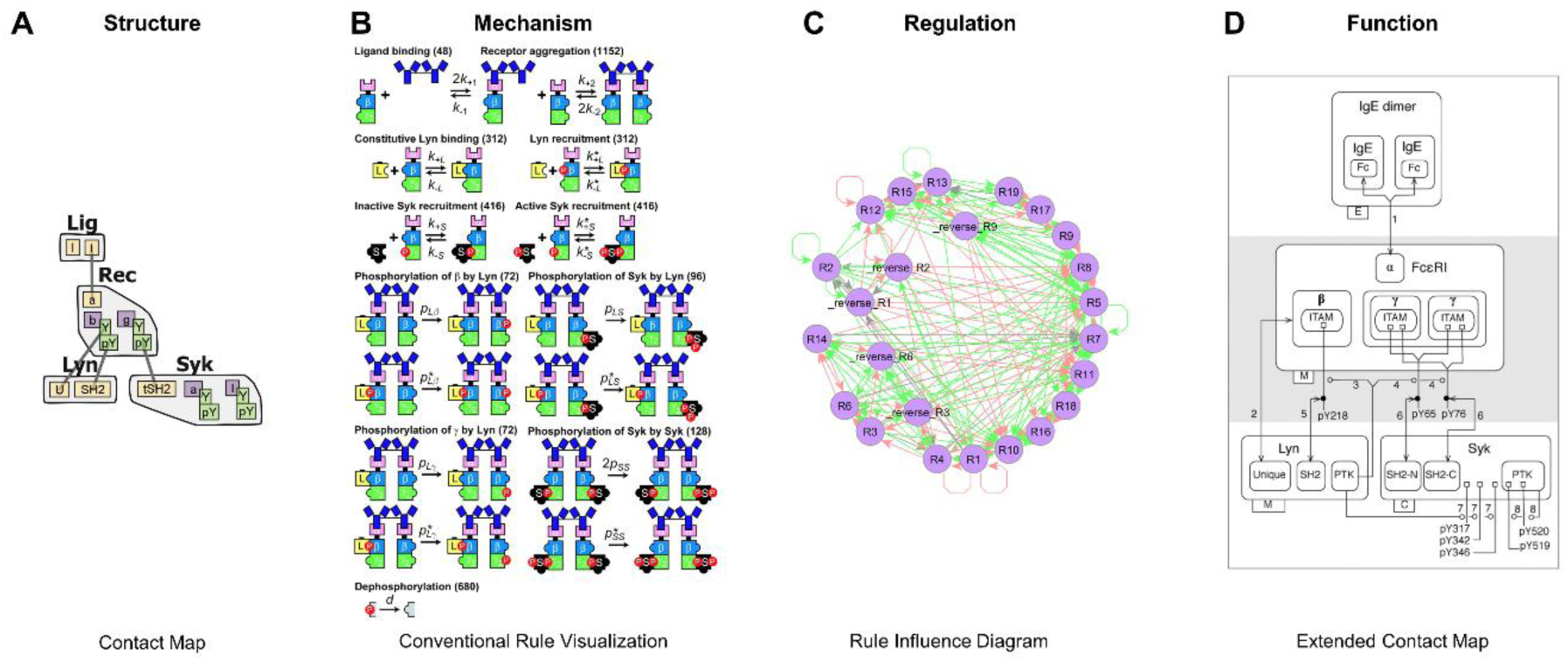
Motivation for new visual tools. Visualization of the different layers of the rule-based model from Faeder et al. [13]. **(A) Structure.** The contact map shows one instance each of the types of structures in the model. **(B) Mechanism.** By convention, each rule is visualized by drawing reactants and products separately. Left and right sides of the rule have to be compared to identify the action of the rule. **(C) Regulation.** Pairwise overlaps between rules are shown as edges on a rule influence diagram. This does not convey the action of rules on model structures and also results in high edge density. **(D) Function.** The extended contact map is an interpreted summary of the facts of the model superimposed on the contact map, but it has to be manually constructed. There is a need for a unified framework that bridges the layers of the model and enables automated visualization of the functional layer. Images reproduced with permission from: (B) Faeder JR et al. Investigation of early events in FcεRI-mediated signaling using a detailed mathematical model. *J Immunol.* (2003) 170: 3769–81. doi:10.4049/jimmunol.170.7.3769 (C) Chylek LA et al. Guidelines for visualizing and annotating rule-based models. *Mol Biosyst.* (2011) 7: 2779–95. doi:10.1039/c1mb05077j

The *contact map* [16] (Fig 3A) is a summary of the structural layer. It shows a single instance of every distinct type of structure present in the system, i.e., molecule, molecular component (binding sites and phospho-sites), internal state of component (unphosphorylated and phosphorylated states) and binding interaction. However, it does not show signal flow between structures. *Conventional rule visualization* (Fig 3B) shows the mechanistic layer by visualizing each rule separately, with reactants and products within each rule drawn separately. Inferring the action of a rule requires comparing reactant and product patterns, whereas inferring the influence between two rules requires comparing patterns across rules. These visual tasks are hindered by the complexity of the pattern graphs that compose the rules and are not scalable to large numbers of rules. The *rule influence diagram* [17] (Fig 3C) is a representation of the regulatory layer. Each node represents a rule and each edge represents an interaction between a pair of rules. Inferring the diagram requires explicitly comparing each pair of rules in the model, which is computationally inefficient. The diagram is also simplistic because it does not show the effect of rules on model structures, and too dense to be useful for visual analysis because of the large number of pairwise overlaps. The *Extended Contact Map* [14] (Fig 3D) is a representation of the functional layer and is a global summary of the model. To build the map, each structure, rule and rule influence is manually interpreted and superimposed on the contact map using ECM conventions [14]. Although it is visually accessible to a wide audience, the ECM is difficult to automate because of the degree of manual interpretation involved. Note that the four layers of information are connected to each other in specific ways that involve comparison and coarse-graining operations. While it is possible to manually delineate these connections for individual models, the state of the art does not provide a systematic approach that can be more generally applied.

In this work, we have produced three major advances over the state of the art for visualization of rule-based models. First, we have developed a novel *compact rule visualization* that does not require left-to-right visual comparison to convey the action of the rule. Second, we have enabled automatic generation of the *rule-derived regulatory graph*, which shows regulatory interactions between rules and model structures, and we show how it can be derived from rules without pairwise comparisons. The regulatory graph was first introduced in the *rxncon* modeling framework [22], which uses a restricted model specification (see Discussion). Here, we show how to infer the graph from the more general rule-based specification, which makes it applicable to a more comprehensive set of models in the widely used BioNetGen [12,23,24], Kappa [16,25] and Simmune [19,26,27] frameworks. Third, we have developed approaches to formally reduce regulatory complexity using user annotation and graph analysis. This makes the rule-derived regulatory graph visualization scalable to models with a large number of rules and produces compact diagrams of the signaling architecture of the system. In the Methods section, we provide the graph definitions and algorithms that bridge the different layers of the model and enable these visualizations. These methods have been implemented in BioNetGen, but will be applicable to Kappa and Simmune models through the proposed interchange format SBML-Multi [28] or the PySB modeling framework [56]. In the Results section, we apply these tools to rule sets of different sizes, from a few rules to hundreds of rules. We also show a comparison of complexity statistics for different visualization methods applied to 27 rule-based models from the literature (listed in S6 Appendix). In the Discussion section, we provide a detailed comparison with other tools in the literature, as well as new directions in which our work can be extended.

## Methods

### Approach

The goal of this work is to formally derive useful visualizations of rule-based models. The input for these methods include BioNetGen patterns and rules, whose formal definitions have been provided in previous literature (see Supplement in [11]). These are converted to intermediary graphs, which are graph data structures defined in this work. Diagrams are generated by applying rendering conventions to the nodes and edges of the intermediary graphs. Here, we briefly discuss intermediary graphs and algorithms for their interconversion and rendering. We provide a detailed specification of algorithms in S7 Appendix and rendering conventions in S8 Appendix respectively.

### Intermediary Graph Types

Intermediary graphs are of two broad types: structure graphs and network graphs (Fig 4A). **Structure graph** denotes a node-labeled undirected graph, with node attributes including but not limited to { NodeIndex, NodeLabel, NodeType }. Multiple nodes with the same NodeLabel and NodeType are allowed, but not with the same NodeIndex. Structure graphs are used to represent objects with predominantly structural information (Fig 4A). **Network graph** denotes an edge-labeled node-labeled bipartite graph. Node attributes are { NodeLabel, NodeType } with NodeLabel being used to uniquely identify nodes. Network graphs are allowed to have only two node types with one node type referring to a structure and the other to a process. Edges are bipartite, i.e. they can only be from a structure node to a process node and not between the same node types. Edges on the network graph have one or more binary attributes that are typically mapped to visual attributes such as color and direction. Network graphs are used to represent the interaction of processes with structures (Fig 4A). Structure graphs and network graphs have precedent in hierarchical graphs [29] and *rxncon* regulatory graphs [22] respectively.

**Fig 4.**
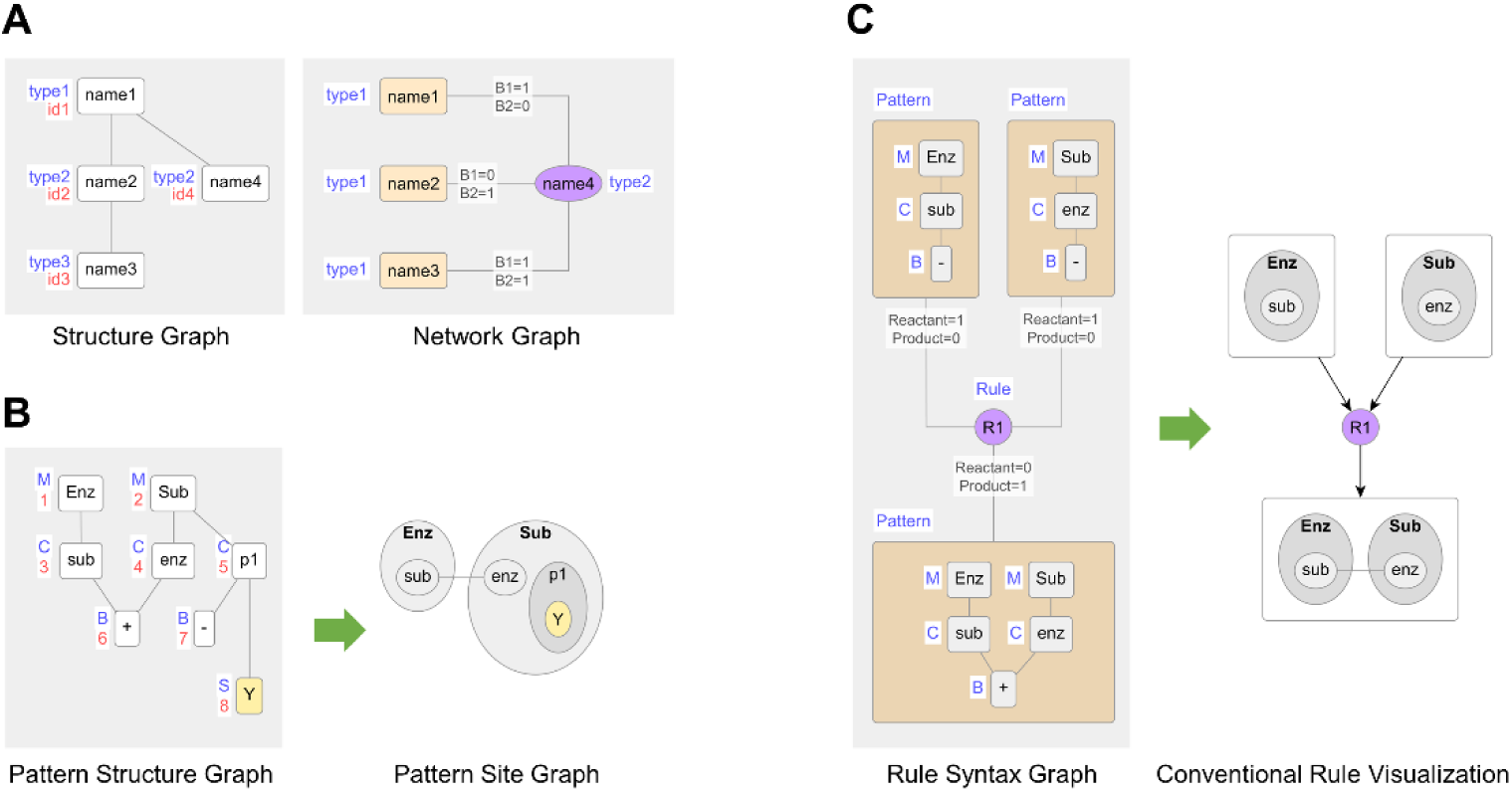
Intermediary graphs and conventional visualizations. **(A)** Structure graphs ({NodeIndex, NodeLabel, NodeType} as node attributes) and network graphs ({NodeLabel, NodeType} as node attributes) are the two types of graph data structures used in this work. Edges on the network graph are bipartite and have one or more edge attributes that can take binary values. **(B)** Patterns are represented by the pattern structure graph, which has molecule, component, binding state and internal state as values for NodeType (denoted M, C, B, S respectively). To visualize a pattern as a site graph, hierarchically nest molecules (nodes 1,2), components (3,4) and internal states (8), show bonds (6) as edges between components, and ignore unbound states (7) **(C)** The rule as specified in the model is represented by the rule syntax graph, a network graph in which one node type embeds pattern structure graphs and the other is labeled with a rule name. To produce a conventional rule visualization, render embedded patterns as site graphs and use edge direction to show reactant and product relationships.

### Visualizing Structure

The **pattern** is the BioNetGen object that is used to specify molecular structures. It is built from molecules, components of molecules, internal states of components and bonds between components [10–12]. Here, we represent the pattern as the **pattern structure graph,** in which each molecule, component, internal state and bond state (bond or unbound state) is represented as a node (Fig 4B, S7.1 in S7 Appendix) ([11,29,30] have similar graph formalisms). To visualize the pattern as a **site graph** (Fig 4B), nest molecules, components and internal states hierarchically and show bonds as edges between components (S8.1 in S8 Appendix). The chemical species and the contact map are both special cases of the pattern structure graph and have the same rendering conventions.

### Visualizing Mechanism

The **reaction rule** is the BioNetGen object that is used to specify kinetic processes. It contains patterns specified as reactants and products [10–12]. The **rule syntax graph** is a network graph in which one type of node embeds pattern structure graphs, the other type of node is labeled with a rule name, and edges between the nodes are typed as reactant or product (Fig 4C). This encodes the reaction rule as it is specified in the model. To achieve a **conventional rule visualization** (Fig 4C), render the embedded patterns as site graphs and use edge direction to show whether the relationship is reactant or product (pattern to rule = reactant, rule to pattern = product).

In conventional rule visualization, reactants and products are drawn separately, so the structures shared between them are drawn twice. Inferring the action of the rule requires visual comparison of reactants and products to identify shared versus unique structures, which does not scale well with pattern complexity. Here, we merge the shared structures between reactant and product sides of the rule to produce a single graph called the **rule structure graph** (Fig 5A, S7.2 and S7.3 in S7 Appendix). The resultant nodes inherit the property of whether they originate from reactant side or product side or both, which is represented as the attribute NodeSide. This explicitly differentiates modified structures from unmodified ones. To obtain a **compact rule visualization**, render unmodified nodes on the rule structure graph using site graph conventions and use graph operation nodes to emphasize the modified structures (Fig 5A, S8.2 in S8 Appendix). Five types of graph operations are permitted in BioNetGen rules (Fig 5B): adding and removing bonds (AddBond/DeleteBond), creating and destroying whole molecules (AddMol, DeleteMol) and changing the internal state label of a component (ChangeState). A rule may have more than one graph operation occurring simultaneously.

**Fig 5.**
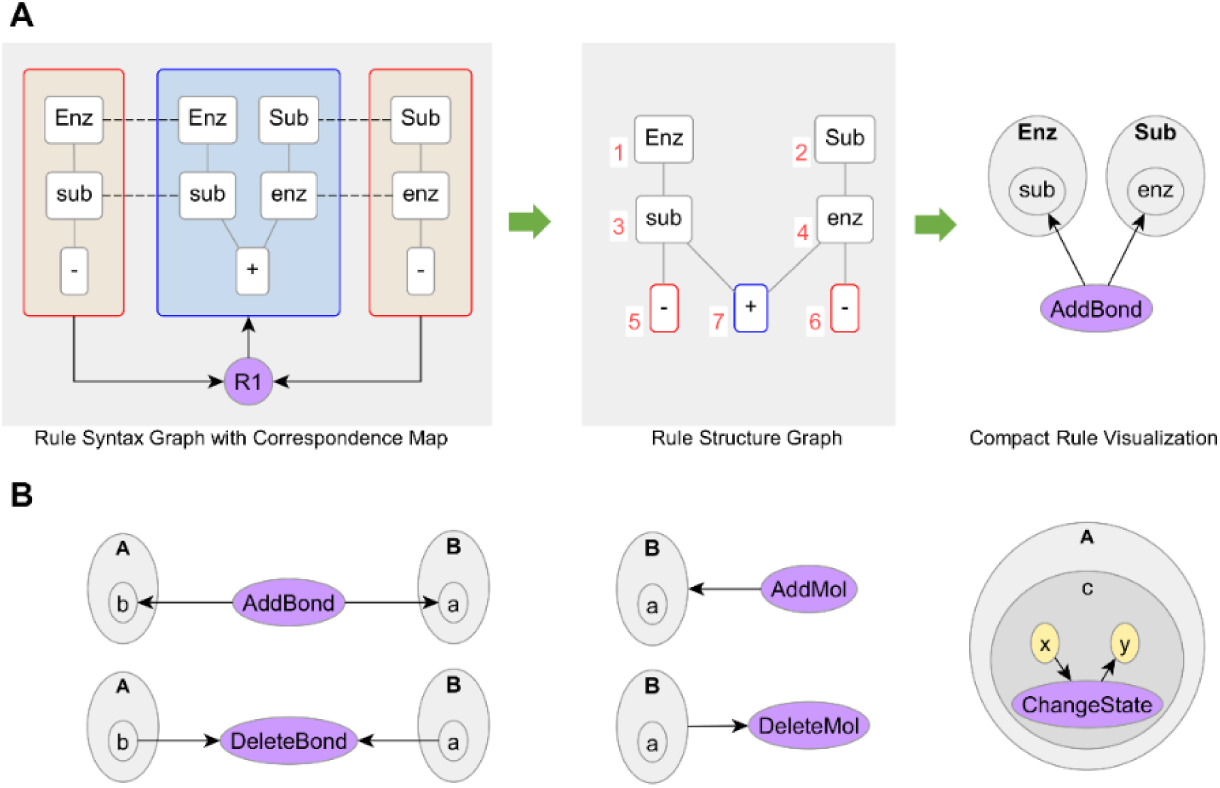
Compact rule visualization. **(A)** Given a rule syntax graph, BioNetGen maps out the shared structures between reactants and products by building a correspondence map (dashed lines). We use the correspondence map to merge reactants and products into a single rule structure graph. Nodes on this graph are also labeled with a side of origin: reactant only (red node border), product only (blue) or both (gray). To visualize rules compactly, render the rule structure graph as a site graph (nodes 1-6) and use graph operation nodes (AddBond) to emphasize the modified structures (node 7). **(B)** BioNetGen has five basic graph operations: adding and removing bonds, creating and deleting molecules and changing state labels of molecular sites. A rule may have one or more instances of these graph operations.

### Visualizing Regulation

Compact rule visualization does not show relationships across rules. To do so, we introduce the notion of **atomic pattern**, a special type of pattern that specifies only a single distinct type of modifiable structure. Atomic patterns include free binding sites and bonds (modifiable by AddBond/DeleteBond), types of molecules (modifiable by AddMol/DeleteMol) and internal states (modifiable by ChangeState). Atomic patterns are determined by examining specific source nodes on rule structure graphs (Fig 6A, S7.4 in S7 Appendix). Since each node on the rule structure graph has a NodeSide attribute (reactant/product/both), this is translated as a relationship between the matched atomic pattern at that node and the rule itself (reactant/product/context). These relationships are represented as binary labels on bipartite edges between the rule and the atomic patterns, resulting in the network graph called the **rule-derived regulatory graph** (Fig 6A, S7.4 in S7 Appendix). To visualize regulatory graphs (Fig 6A, S8.3 in S8 Appendix), we apply distinct visual attributes to the two node types (atomic pattern, rule) and the three edge types (reactant, product, context). Individual rule regulatory graphs are merged and processed to build the **model regulatory graph** (Fig 6B, S7.5 in S7 Appendix) on which the influence between any two rules can be traced as a path through an atomic pattern. The rules in a model draw from the same finite pool of structure types and binding interactions, so the model regulatory graph grows primarily with the number of rules in the model.

**Fig 6.**
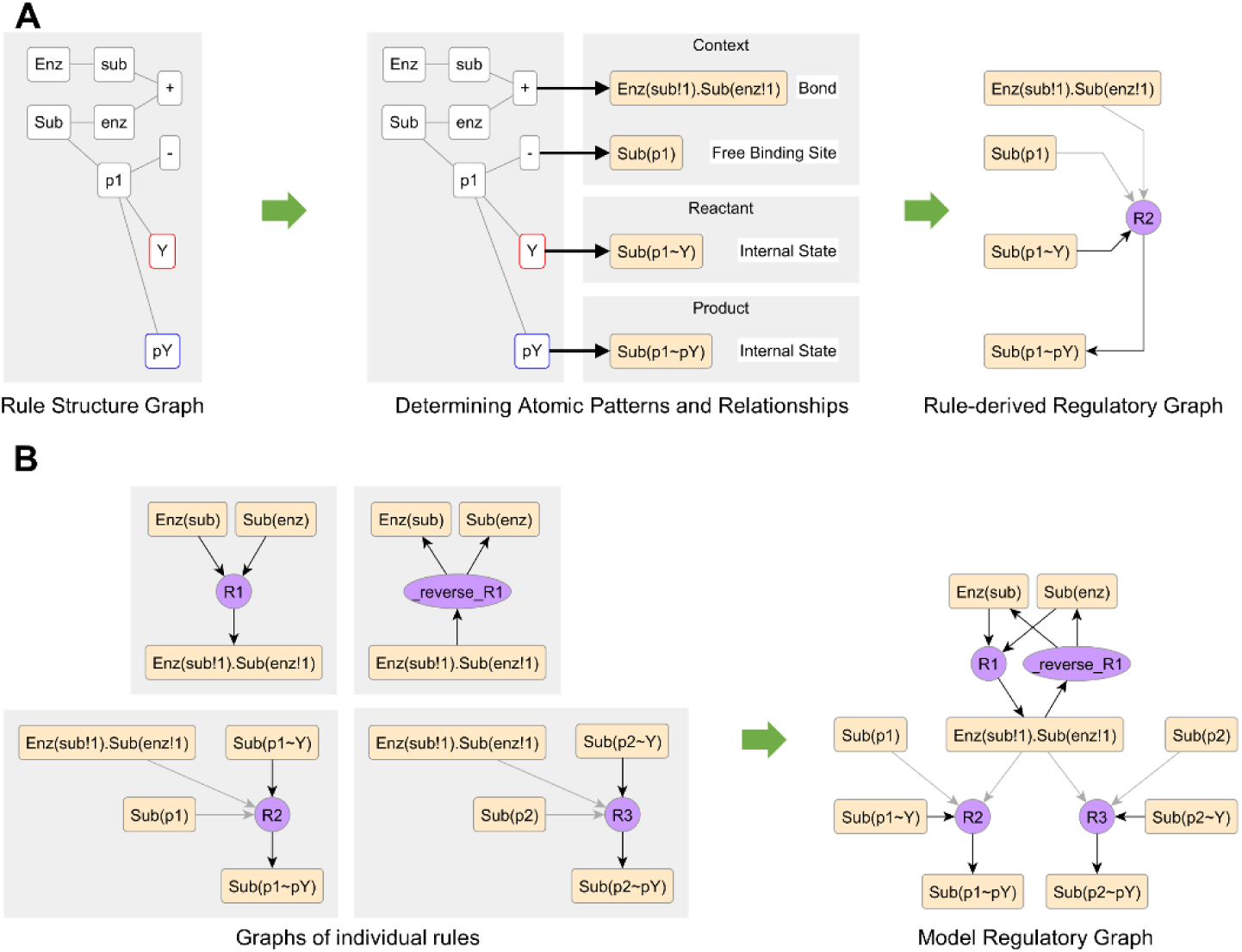
Rule-derived regulatory graph. **(A)** An atomic pattern represents a distinct type of modifiable structure: a free binding site, a bond, an internal state or a molecule type. It has a name specified using pattern syntax (brackets = containership, !{tag}=bond, ~{label}=internal state). Atomic patterns are identified by examining source nodes on the rule structure graph (bold arrows). The rule-derived regulatory graph is a network graph with edges between atomic patterns and a node labeled with the rule name. The NodeSide attribute of the source node (border color red=left, blue=right, gray=both) determines edge attributes on the regulatory graph (reactant, product and/or context). Context edges are rendered with a light color and reactant/product edges have a dark color. **(B)** Aggregating regulatory graphs of individual rules results in the model regulatory graph. The model shown here is an enzyme reversibly binding to substrate (rules R1 and _reverse_R1) and phosphorylating two sites on the substrate (rules R2 and R3 respectively).

### Visualizing Function

The model regulatory graph provides an explicit description of signal flow that is formally derived from the rules (Fig 7A), but this information needs to be simplified in order to build compact pathway diagrams. This involves four steps: background removal, grouping atomic patterns, grouping rules and compressing groups. Background removal consists of removing atomic patterns and rules that are redundant for model comprehension, e.g., free binding sites and unphosphorylated states (Fig. 7B, S7.6 in S7 Appendix). This greatly reduces the size of the graph. Grouping atomic patterns involves identifying similarity between sites in the system, e.g., phospho-sites on the same molecule or bonds between the same pair of molecules (Fig 7C). Grouping rules requires computing similarity between rules based on their edge signatures, i.e., based on how each rule interacts with groups of atomic patterns (Fig 7D, S7.7 and S7.8 in S7 Appendix). Compressing groups involves replacing groups of nodes by a representative group node and combining edges incident on individual members of the group and remapping them to the representative group node (Fig 7E, S7.9 in S7 Appendix). Background removal and atomic pattern grouping are performed using default heuristics whose choices can be modified with user input, then rule grouping is performed automatically based on those settings. The edge signature can be strict or permissive, i.e. it considers all three edge types or it considers only reactant and product edges and ignores context edges respectively. Using a permissive edge signature results in identifying a broader grouping of rules. The choice of edge signature as well as the modifiers to default heuristics enable customizing the output graph to account for nuances of individual models, the target audience, and the desired level of compression.

**Fig 7.**
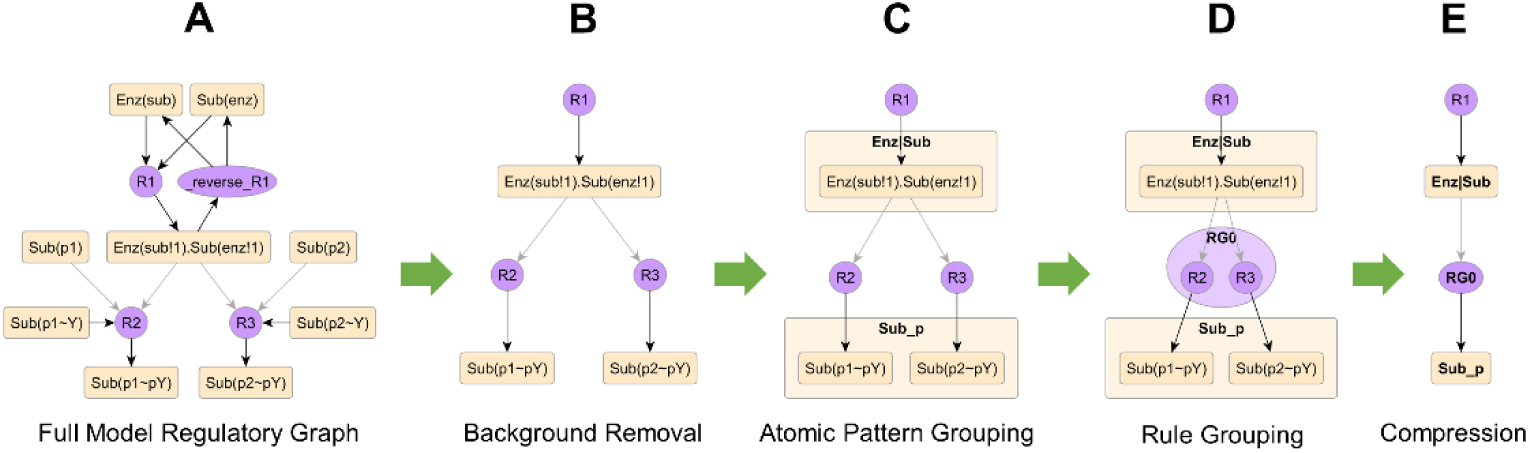
Complexity reduction approaches. **(A)** The full model regulatory graph from Fig 6. **(B)** Nodes redundant for human comprehension are tagged as background and removed. Here, that includes free binding sites and the unbinding rule. **(C)** Atomic patterns are classified into groups. Here, the groups used are Enz|Sub to denote the enzyme-substrate bond and Sub_p to denote the phosphorylated sites. Steps (B) and (C) can be performed by a heuristic that can be modified with user annotation. **(D)** Rules are classified into groups by an automated comparison of their edge signatures. Here, rules R2 and R3 have similar edge signatures: a context edge from Enz|Sub and a product edge into Sub_p, so they are grouped under the same label RG0. **(E)** Each group of nodes is replaced by a single representative node and edges incident on individual members of the group are merged together.

### Algorithm Time Complexity

If *p* represents the size of a pattern, building a pattern structure graph from the pattern syntax is *O(p)* (S7.1 in S7 Appendix). If *mL, mR* represent the size of patterns on left and right sides of a rule respectively, building a correspondence map between reactants and products before merging is *O(mL*mR)* (S7.2 in S7 Appendix). Building the rule structure graph from the correspondence map (S7.3 in S7 Appendix) and building the rule regulatory graph from the rule structure graph (S7.4 in S7 Appendix) are both *O(m)* where *m* represents the size of the rule. Rules are typically finite and bounded in size (shown for 2239 rules in Fig S1), so the contribution of rule size can be assumed to be *O(1)*. Given this, building the regulatory graph (S7.5 in S7 Appendix), removing background (S7.6 in in S7 Appendix), identifying rule groups using edge signatures (S7.7 and S7.8 in S7 Appendix) and collapsing group nodes (S7.9 in S7 Appendix), are all *O(n)*, where *n* is the number of rules.

### Implementation

The methods described here have been integrated into the BioNetGen framework [12,23,24] and are freely available as part of the open source BioNetGen distribution at http://bionetgen.org. A typical visualization procedure involves calling a “visualize()” method from BioNetGen, which outputs a file in GML format (graph modeling language) [31,32], which is then imported and laid out using a dedicated graph visualization tool. The graphs in this work were laid out using yEd, a commercial graph editor freely available at http://yworks.com/yed. User input for background removal and atomic pattern grouping heuristics is provided using text-based template files. A template file with the default choices can be automatically generated for each model and then modified by the user.

### Graph Complexity

We compiled a list of 27 rule-based models from the literature containing a total of 2239 rules (S6 Appendix). The number of rules per model ranged from 6 to 625. To assess rule size, we computed number of nodes for rule syntax graphs, rule structure graphs and rule-derived regulatory graphs for each rule. To compare visual complexity of different model visualization tools, we applied those tools to the 27 models and compared number of nodes and number of edges per node, counting hierarchical relationships between nodes also as edges. The visualization methods applied are conventional rule visualization, compact rule visualization, Simmune Network Viewer, contact map, rule influence diagram and four types of regulatory graphs: full model regulatory graphs, regulatory graphs with background removed, compressed regulatory graphs using a strict edge signature, and compressed regulatory graphs with a permissive edge signature. Background selection and classification of atomic patterns into groups were performed using default heuristics. To compare rule-based models and reaction networks, we generated equivalent reaction networks for 20 of the 27 rule-based models using methods described previously [12]. Then we counted species and reactions in reaction networks and patterns and reaction rules in rule-based models respectively. For the remaining 7 models, it was not possible to generate a finite-sized network using available computational resources.

## Results

### Visualizing Interactions of Reaction Rules

Visualizing individual rules promotes understanding the structural and kinetic assumptions encoded in the model. Conventional rule visualization entails drawing reactants and products of the rule separately and inferring the action of the rule by visual left-to-right comparison. In contrast, compact rule visualization shows the action of the rule explicitly using nodes that represent operations such as adding or removing bonds, creating or deleting molecules and changing internal states of sites. We demonstrate compact rule visualization using four rules from Faeder et al. [13] that model the interaction of Lyn kinase with the FcεRI receptor. Fig 8A shows two rules R3 and R6 that model binding of Lyn to receptor. Rule R3 models constitutive recruitment: the binding site on the receptor is unphosphorylated and the binding domain on Lyn is the unique domain U. R6 models active recruitment: the binding site on the receptor is phosphorylated and the binding domain on Lyn is the SH2 domain. Fig 8B shows two rules that model phosphorylation of receptor by Lyn. The phosphorylation occurs across the dimer interface: Lyn recruited to one side of the dimer phosphorylates a site on the other side of the dimer. In R4, the Lyn kinase is constitutively recruited and in R7, it is actively recruited.

**Fig 8.**
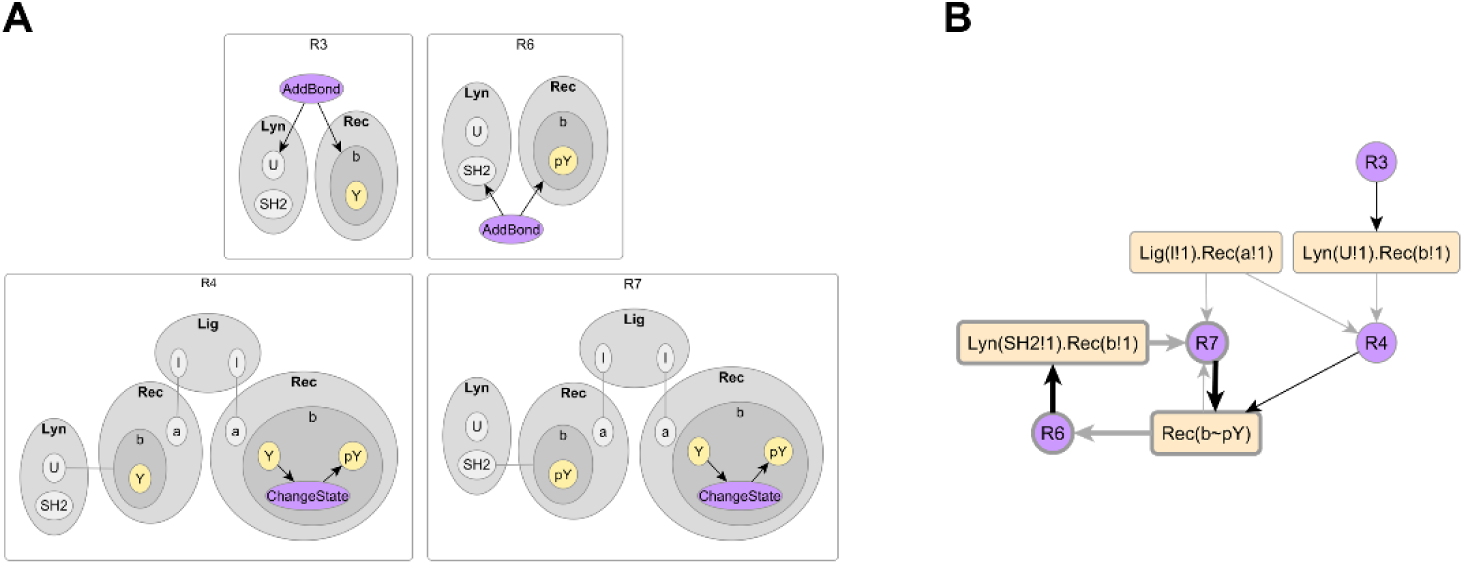
Lyn-FcεRI interactions. **(A) Lyn-FcεRI binding.** Rules R3 and R6 model constitutive and activated binding of Lyn kinase to β site of receptor respectively. The β site is unphosphorylated in R3 and phosphorylated in R6. The binding domain of Lyn is U in R3 and SH2 in R6. **(B) Phosphorylation of FcεRI.** Rules R4 and R7 model trans-phosphorylation of β-site in ligand-crosslinked receptor dimer. The kinase is constitutively recruited Lyn in R4 and actively recruited Lyn in R7. **(C) Positive feedback loop.** The regulatory graph of the four rules (with free binding sites and unphosphorylated states removed) shows the architecture of this system, which includes a positive feedback between activated Lyn recruitment and receptor phosphorylation (highlighted with bold lines). See Fig 6 for information on the naming syntax for sites.

The structures common to each pair of rules mediate the interactions between them. For the four rules shown here, it is possible to infer regulatory interactions by analyzing which structures are modified or required in each rule. R4 requires constitutively bound Lyn, which is produced by R3. R6 requires phosphorylated β site, which is produced by both R4 and R7. R7 requires actively bound Lyn, which is produced by R6. Regulatory motifs emerge from the combined effect of individual overlaps. Here, recruited Lyn (constitutively or actively recruited) promotes receptor phosphorylation, which in turn promotes active Lyn recruitment, and the emergent architecture is a positive feedback loop. In general, the overlap between two rules is some complex combination of instances of structures and prior to this work, the only way to infer regulation from a set of reaction rules was to compare or simulate the effect of each pair of rules [16,17]. For the published rule-based models, the emergent signaling architecture is usually identified and diagrammed by manually interpreting the rules (e.g. [33]).

The rule-derived regulatory graph enables an automated visualization of the regulatory layer without having to analyze pairs of rules. First, the structures specified in each rule are projected into a simpler space of basic structure types, called atomic patterns. Then, the relationship of the rule to each atomic pattern is inferred from the rule as a simple bipartite relationship (reactant/product/context) and is represented as an edge between rule and atomic pattern. The regulatory graphs of individual rules are aggregated into a regulatory graph of the system (Fig 8B) on which overlaps between rules are identifiable as paths mediated by atomic patterns. For example, the overlap between R3 and R4 is via the bond atomic pattern that represents constitutively recruited Lyn (Fig 8B). Emergent signaling motifs can be identified as paths traced on the regulatory graph, such as the positive feedback loop between receptor phosphorylation and active Lyn recruitment highlighted in Fig 8B. Thus, the rule-derived regulatory graph provides an automated visualization of rule-based models that enables rapid identification of network-level motifs.

### Tuning Display of Regulatory Complexity

Applied to common rule-based models, the rule-derived regulatory graph has greater size and complexity than a typical pathway diagram. Here, we show that a simple pathway representation can be achieved by formally compressing the rule-derived regulatory graph using user annotation and graph analysis. We also show, through application to the model of Faeder et al. [13] (Fig 3), that compression is flexible and can be adjusted to account for the nuances of a specific model and for the level of complexity desired in the eventual diagram.

The Faeder et al. model [13] describes activation of kinase Syk by ligand-induced aggregation of FcεRI receptors. As shown in Fig 3A the model has a receptor, a bivalent ligand and two kinases Lyn and Syk that are recruited to the β and γ sites of the receptor respectively. The ligand crosslinks receptors into a dimer and Lyn recruited to one side of the dimer phosphorylates and activates sites on the other side of the dimer that recruit Lyn and Syk. Kinase activity across a dimer interface is a frequently encountered mechanism called *trans-*phosphorylation and is common among receptor tyrosine kinases (RTKs) [34] and non-receptor tyrosine kinases recruited to receptors (nRTKs) [35]. Syk has two phospho-sites, one on its linker region and one on its activation loop, which are *trans*-phosphorylated by recruited Lyn and recruited Syk respectively. Lyn itself may be recruited in two ways: constitutively to an unphosphorylated β site and upon ligand stimulation to a phosphorylated β site. The model has 24 rules and the full rule-derived regulatory graph without any compression is presented in Fig 9A. While this graph shows all regulatory interactions present in the model, it is difficult to understand because of its size and complexity.

**Fig 9.**
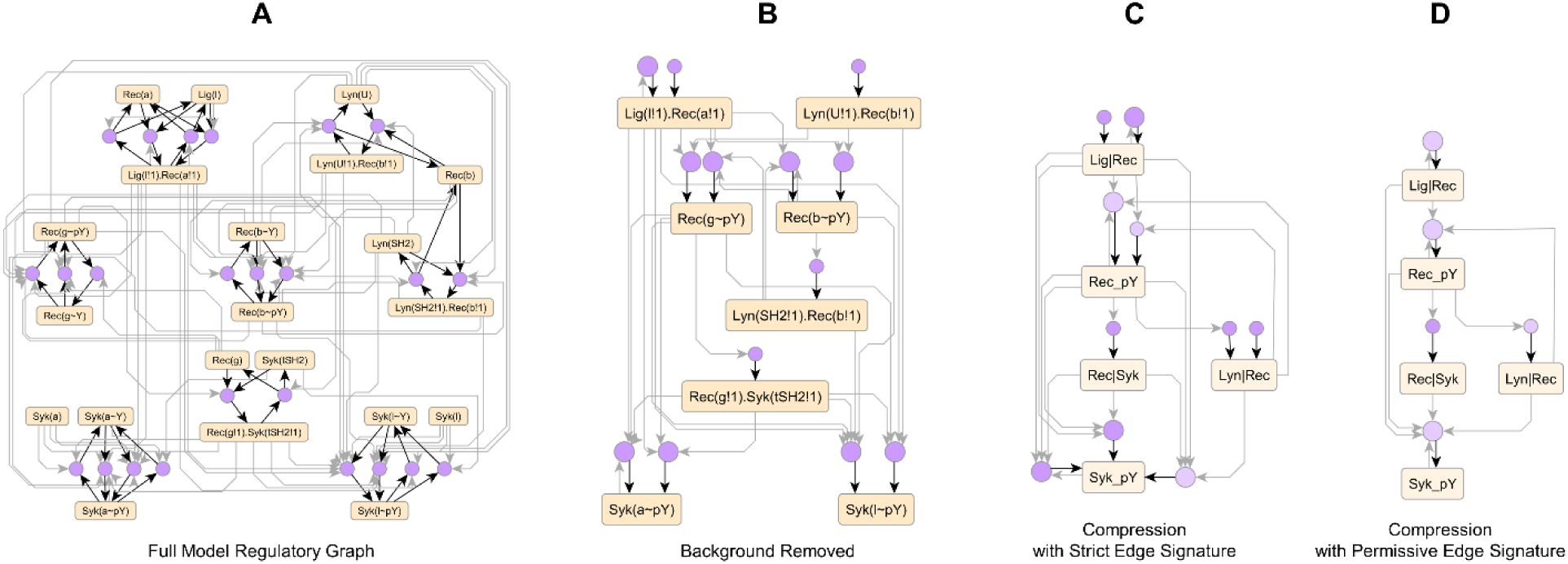
Regulatory complexity of the FcεRI model. Graphs generated from the Faeder et al. model [13] of FcεRI signaling resolve regulatory architecture of the model to different extents. **(A)** The full model regulatory graph resolves each rule and each type of structure in the model. **(B)** Removing background nodes reduces graph size, but still resolves most regulatory interactions. The nodes removed include free binding sites, unphosphorylated states, dissociation rules and dephosphorylation rules. **(C)** Merging groups of structures and rules coarse-grains regulatory complexity. Groups of structures are assigned by a heuristic: groups of phospho-sites on the same molecule (e.g. Rec_pY, Syk_pY) and groups of bonds between the same molecule pair (e.g. Lig|Rec, Rec|Syk and Lyn|Rec). Here, the strict edge signature was used, which resolves variations of the same process that have different context, e.g., the three process nodes with a product relationship to Syk_pY. **(D)** A permissive edge signature merges rules more aggressively and does not resolve contextual variants, e.g., the single node with a product relationship to Syk_pY, resulting in a compact and simplified diagram of the signal architecture. Note that the positive feedback loops between Lyn-receptor binding and receptor phosphorylation and from Syk phosphorylation to itself are preserved through diagrams A-D. For (C) and (D), the grouping prior to compression is shown in Fig S2A and Fig S2B respectively.

The first step in complexity reduction is to identify atomic patterns and rules that are redundant for human comprehension. For example, the unphosphorylated states of sites in the system are rarely considered part of the active signaling architecture when discussing a model. Similarly, if a binding event is considered part of the propagated signal, then the corresponding dissociation process is typically ignored as a background process. Here, we provide a default heuristic that removes unbound states, default states, and immediate reverses of rules, but these choices can be modified by the user to account for other types of regulation. For the Faeder et al. model, in addition to the default choices, we added dephosphorylation rules to the background also. The resultant graph in Fig 9B is less than half the size of the full graph but still shows all of the relevant regulatory interactions.

The second step in complexity reduction is to group similar structures on the regulatory graph. We provide a default heuristic that groups identical state labels present on components of the same molecule and bonds between the same pair of molecules, but these can be changed to account for other meaningful ways in which structures can be organized into groups. Following background removal, the default grouping heuristic results in the atomic pattern groups Rec_pY, Syk_pY, Lig|Rec, Rec|Syk and Lyn|Rec (Fig 9C). The Rec_pY group consists of phosphorylated states of β and γ sites of receptor and the Syk_pY group consists of phosphorylated states of activation loop and linker region of Syk respectively. The Lig|Rec and Rec|Syk groups consist of the ligand-receptor and receptor-Syk bonds respectively and the Rec|Lyn group consists of the two receptor-Lyn bond types.

The third step in complexity reduction is to group rules based on whether they interact with the same types of structures in the same way. Rules are placed into the same group if they share the same *edge signature*, which is computed from the labels of adjacent edges and nodes and also takes into account the grouping of atomic patterns. We have enabled two types of edge signatures: strict and permissive. The strict edge signature considers reactant, product and context edges, whereas the permissive edge signature ignores context edges. Application of the rule-grouping algorithm to the regulatory graph in Fig 9B results in the graph shown in Fig S2A using the strict edge signature and the graph in Fig S2B using the permissive edge signature. Rule groups in Fig S2B are larger than in Fig S2A because of the relaxed criterion for computing groups.

The fourth step in complexity reduction is to compress the graph by replacing each group of nodes with a single representative node labeled with the group name. Edges on individual members of a group are merged onto the node that represents the group. Compressing the graphs in Figs S2A and S2B resulted in the graphs in Figs 9C and Fig 9D respectively. The choice of background, atomic pattern groups and edge signature tunes the degree to which individual regulatory interactions are resolved on the compressed graph. For example, Fig 9C resolves three contextual variants of Syk phosphorylation, namely one that is Lyn-dependent and two that are independent of Lyn, of which one has feedback from phosphorylated Syk and one has no feedback. On the other hand, Fig 9D uses a single node to represent Syk phosphorylation, resulting in a more simplistic picture that does not resolve contextual variants. Either Fig 9C or 9D can be used as a pathway diagram to communicate the signal architecture of the model, highlighting the feedback between Lyn recruitment and receptor phosphorylation, the dependence of Syk phosphorylation on Lyn, and the additional feedback from phosphorylated Syk.

### Visualizing Reaction Rule Libraries

A number of recently published rule-based models have focused on building catalogues of kinetic interactions downstream from different receptor types [6–9]. The process of grouping and compressing the regulatory graph is scalable to large numbers of rules, which enables visualization of such libraries. Consider the FcεRI library of rules constructed by Chylek et al. [7] and the signaling model of the ErbB receptor family constructed by Creamer et al. [9]. The FcεRI library has 17 molecule types and 178 reaction rules and the ErbB library has 19 molecule types and 625 reaction rules. For both systems, detailed extended contact maps (ECMs) [14] were published along with the model to communicate its architecture. These maps were constructed manually and provided an interpretation of the model rather than a formal representation. Here, we show that it is possible to derive an informative description of signaling from the model itself by applying complexity reduction methods to the rule-derived regulatory graph. Figs 10 and 11 show compressed regulatory graphs of the FcεRI and ErbB models respectively, and these are the first automatically generated visualizations for these libraries that show signal flow.

**Fig 10.**
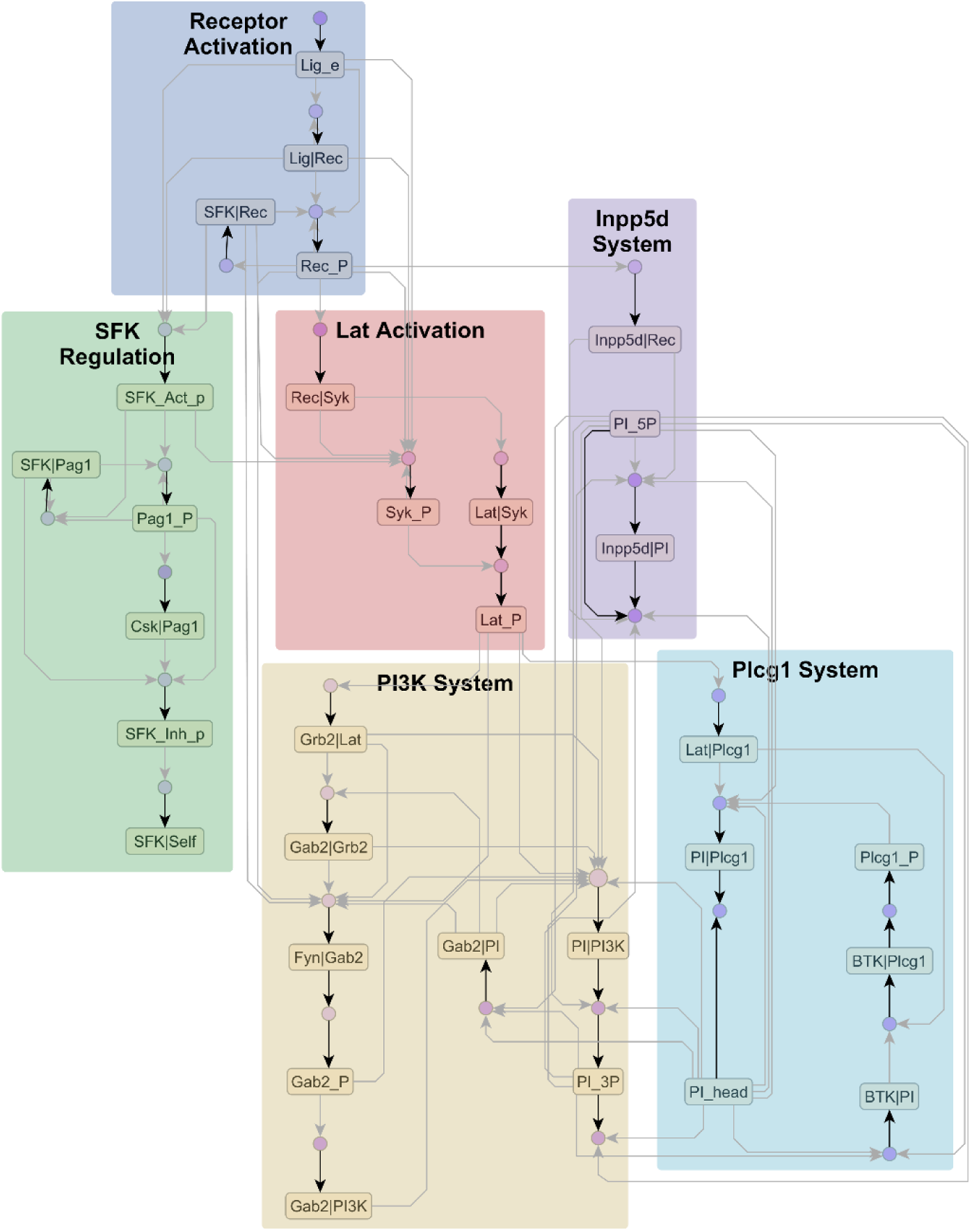
FcεRI library of rules. Compressed regulatory graph (63 nodes, 112 edges) generated from the FcεRI model of Chylek et al. [7] with 178 rules. The uncompressed graph has 305 nodes and 1076 edges. The model elements can be roughly classified into six subsystems shown above.

**Fig 11.**
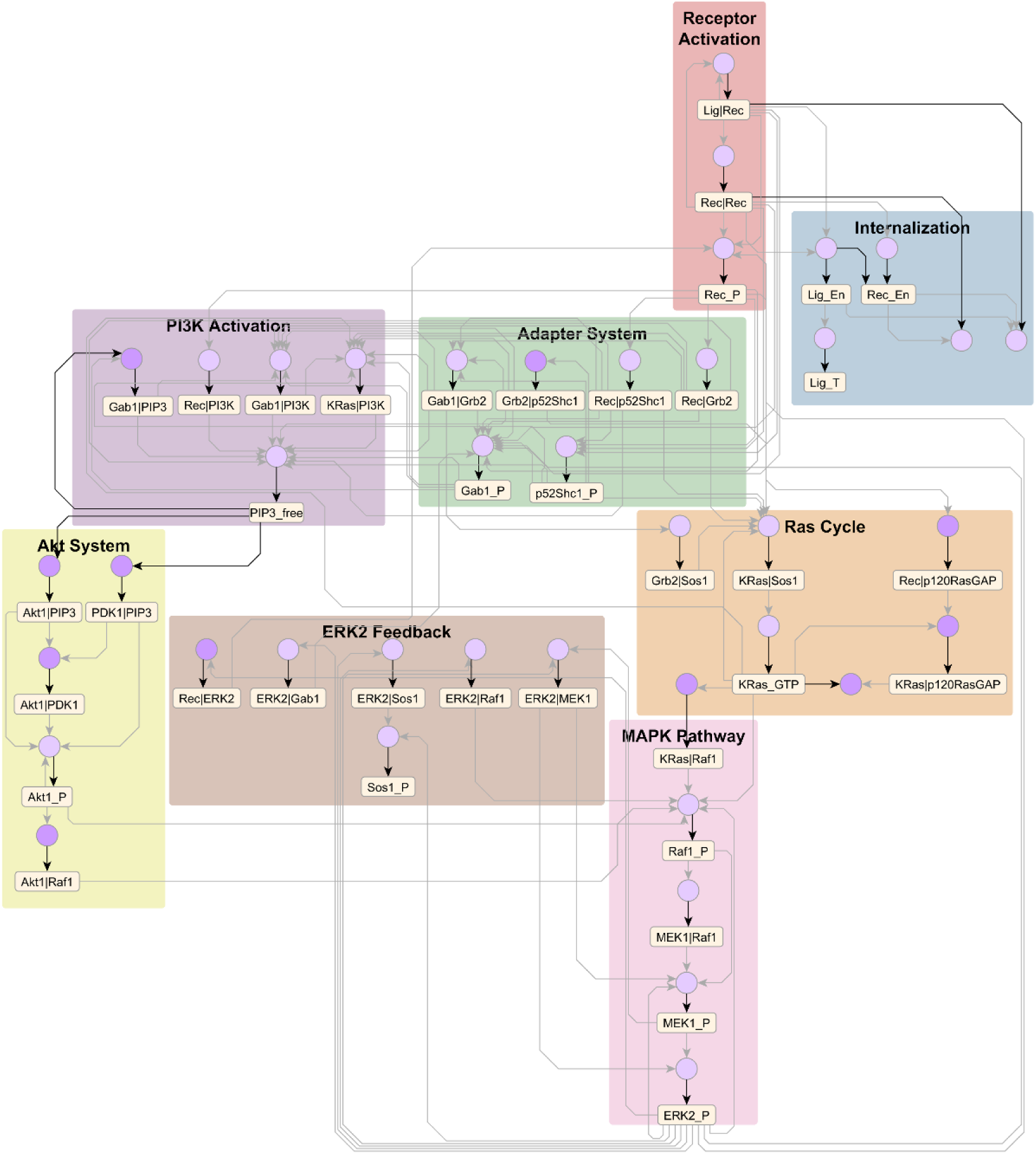
ErbB library of rules. Compressed regulatory graph (79 nodes, 144 edges) generated from the ErbB model of Creamer et al. [9]) with 625 rules. The uncompressed graph has 930 nodes and 5269 edges. The model elements can be roughly classified into eight subsystems shown above.

To generate the compressed regulatory graphs, the default heuristic for background selection and grouping was used with a few user modifications. The nodes removed by default include free binding sites, unphosphorylated states and dissociation rules. With user input, background phosphorylation and dephosphorylation rules as well as constitutive processes were removed. By default, phosphorylation sites on each molecule were grouped together as were binding interactions between the same pair of molecules. However, when functionally similar molecules are present in the system, we found that it was useful to group analogous sites across molecule types rather than dissimilar sites within the same molecule. In the FcεRI model, Lyn and Fyn are Src family kinases that have similar functional roles and are regulated similarly, so analogous phospho-sites and bonds on Lyn and Fyn were grouped together. As a result, on the FcεRI graph (Fig 10), regulatory interactions are mapped to a generic Src family kinase, denoted SFK. In the ErbB model, a similar approach was deployed for the four receptor types (EGFR, ErbB2, ErbB3, ErbB4 grouped under Rec) and the two ligand types (EGF, HRG grouped under Lig) by grouping similar sites across molecules. The four receptor types can bind each other in 10 different ways to form dimers, all of which were grouped under a single heading Rec|Rec. These choices resulted in a dramatic reduction of complexity, resulting in a compact graph that shows signaling from a generic receptor of the ErbB family binding a ligand and forming a dimer (Fig 11). Note that these grouping choices were made for demonstration, so it is possible to make different grouping choices and resolve functionality at a finer level, e.g. by grouping Lyn and Fyn sites separately, or grouping EGFR/ErbB2 phospho-sites separately from ErbB3/ErbB4 phospho-sites.

A major advantage of the regulatory graph is that network-level motifs can be identified by graph analysis. This is very useful for models where the interactions between individual rules are not immediately obvious due to the size of the rule set or the complexity of encoded rules. In Fig 12, we show some of the signaling motifs that emerge from the interactions encoded in the FcεRI library. Kinases Lyn and Fyn, collectively called SFKs, are recruited to sites on the receptor as well as on Pag1 and these binding interactions are enhanced by phosphorylation of those sites by SFKs to form a positive feedback loop (Fig 12A). Similarly, phosphorylation of Syk by Syk leads to increased Syk phosphorylation in a positive feedback loop (Fig 12A). On the SFKs, phospho-sites are classified into positive and negative regulators, i.e. activating sites and inhibitory sites respectively. Auto-regulation is observed when the inhibitory sites are phosphorylated by Csk in a cascade promoted by the activating sites (Fig 12B). Phosphorylated Lat enables Plcg1 phosphorylation via two converging branches of interactions that constitute a coherent feed-forward loop. On one hand, Plcg1 is recruited directly to phosphorylated Lat. On the other hand, the Plcg1-modifying kinase BTK is recruited to PIP3, which is produced by activated PI3K, which in turn is enabled by phosphorylated Lat (Fig 12C). Note that PIP3 was treated as a structured molecule, so there are separate atomic patterns representing the 3’ and 5’ phosphate moieties on the phosphoinositide molecule, i.e., PI_3P and PI_5P respectively. In Fig 12D, we see that there are two branches initiated from phosphorylated receptor with opposing effects on PIP3, which occurs when PI is phosphorylated at both the 3’ (PI_3P) and 5’ (PI_5P) sites. PI3K activity produces PIP3 by adding 3’-phosphate and Inpp5d consumes PIP3 by removing 5’-phosphate. The two branches constitute an incoherent feed forward loop. In the original paper by Chylek et al. (2014), these motifs were identified by manually interpreting the rules, but here they are clearly identifiable on the model regulatory graph.

**Fig 12.**
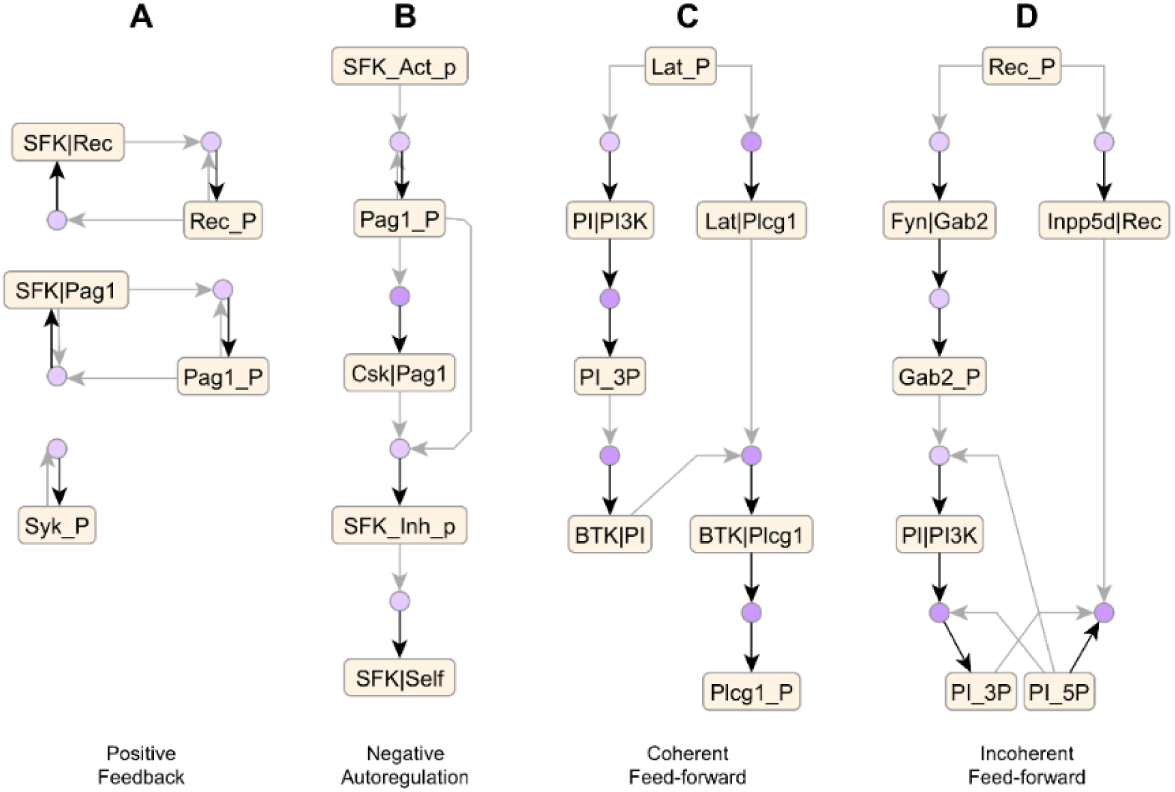
Signaling motifs in FcεRI library. Analysis of the regulatory graph of the FcεRI library in Fig. 10 reveals the regulatory motifs in the system. **(A)** Positive feedback loops enhance binding of Src family kinases Lyn and Fyn (denoted SFK) to receptor and Pag1 scaffold as well as Syk phosphorylation by Syk. **(B)** Pag1 phosphorylated by SFKs recruits Csk, which negatively regulates SFKs by phosphorylation. **(C)** A coherent feed-forward loop activates Plcg1 from phosphorylated Lat. **(D)** An incoherent feed-forward loop involving enzymes PI3K and Inpp5d regulates levels of phosphoinositide PIP3, which is phosphorylated at both 3’ and 5’ hydroxyl positions (denoted PI_3P and PI_5P respectively).

### Comparing Complexity of Visualization Tools

The goal of this work was to enable visualizations that are both automated and comprehensible over a wide range of model sizes. To evaluate readability of various visualizations across models, we first built an analysis set of 243 graphs by applying 9 types of visualizations to 27 rule-based models from the literature. The models ranged in size between 6 and 625 rules (S6 Appendix, S9 Dataset). Next, we needed suitable metrics to evaluate readability. In a study by Ghoniem et al [36], user performance on various graph analysis tasks was found to decrease with increasing graph size *n* and edge density 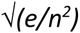, where *n* and *e* refer to the number of nodes and edges on the graph respectively. For the graphs in our analysis set, graph size *n* had a sufficient spread (CV=3.39), but the metric 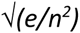 resulted in values with low magnitude and narrow spread (mean=0.094, CV= 1.03). This is because the graphs in the analysis set are at a different regime of edge density than the graphs used in Ghoniem et al., whose 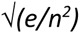 values ranged between 0.2 and 0.6. So we used the metric *e/n* instead to represent edge density, and we found this to be more discriminatory for the graphs in our dataset (CV=2.54). We report the distributions of graph size *n* and edge density *e/n* for each of the 9 types of visualizations in Fig S3. We found some consistent trends in the relative placements of visualizations, which we demonstrate by showing the center of each distribution in Fig 13. The geometric mean was used as a measure of centrality since the distributions varied over few orders of magnitude.

**Fig 13.**
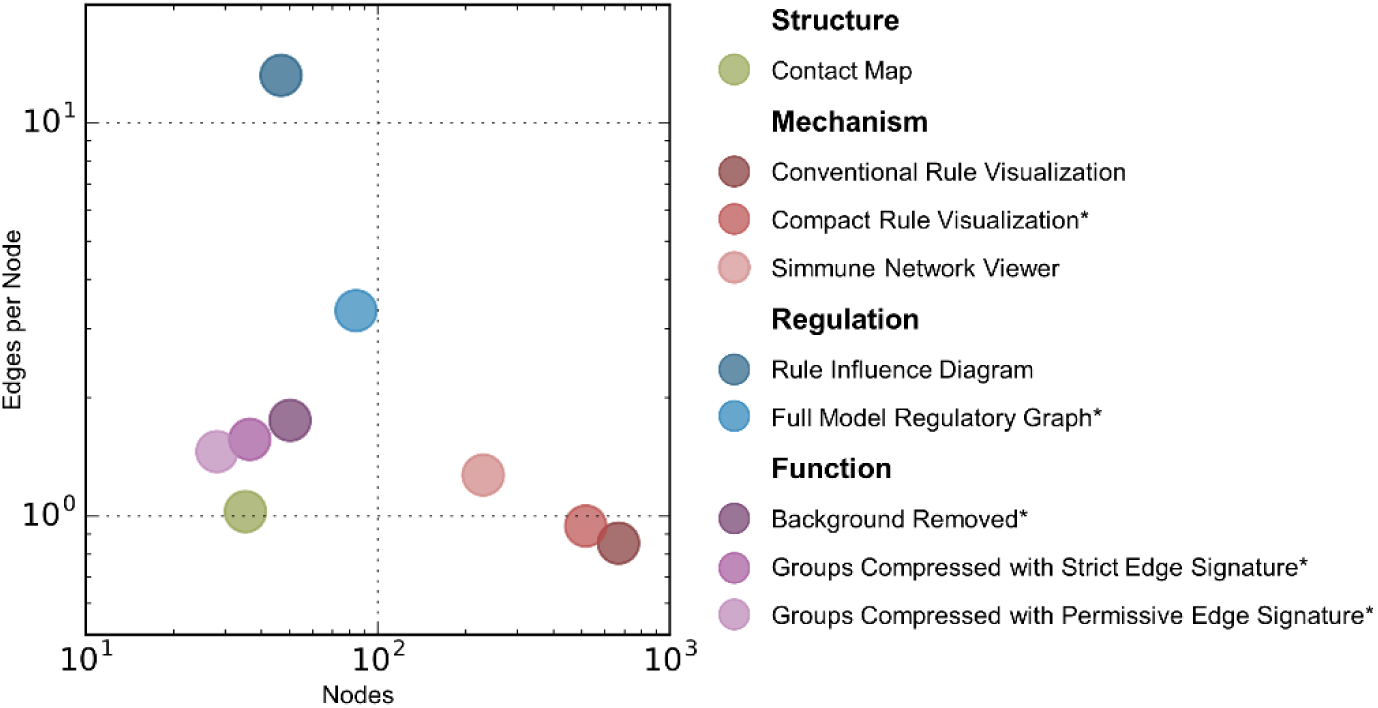
Mean graph sizes for various model visualizations. Each data point shows the geometric mean of graph size and edge density metrics evaluated over 27 rule-based models for each visualization type. The visualization types are: contact map, conventionally visualized rules, compactly visualized rules, Simmune Network Viewer diagram, rule influence diagram, full model regulatory graph, regulatory graph with background removed, regulatory graph compressed with strict edge signature, and regulatory graph compressed with permissive edge signature. Default heuristics were used for background removal and group assignment. Each type of visualization is color-coded based on the layer of information it represents: structure, mechanism, regulation or function. Asterisks (*) indicate the methods developed in this work. Fig S3 shows the full dataset.

The 9 visualizations that are compared in Fig 13 include contact maps, conventional and compact rule visualizations, Simmune Network Viewer diagrams, rule influence diagrams and four types of regulatory graphs: full model regulatory graph, graph with background nodes removed, and compressed graphs generated using either a strict or permissive edge signature. Fig 13 shows that the contact map has on average a small size and low edge density. Conventional and compact rule visualizations have the largest size but low edge density. In comparison, the rule influence diagram is much smaller but has the highest edge density by far. The full model regulatory graph occupies an intermediate position between the rule influence diagram and the rule visualizations on both measures. Background removal drastically lowers both size and density, which can be reduced even more by compressing the graph using strict or permissive edge signatures. The Simmune Network Viewer format and its statistics will be treated in the Discussion section.

The contact map is optimal for visual understanding but only displays the structural layer of the rule-based model, so it is insufficient for understanding regulation. Rule visualizations, which depict the mechanistic layer, have a large number of nodes because rules are composed of complex graphs, but they have low edge density because they do not show regulatory relationships between rules. The rule influence diagram and rule-derived regulatory graphs both show regulatory interactions, and thus convey much more information than contact map or rule visualizations. The rule-derived regulatory graph introduces atomic patterns to mediate relationships between rules, so it has a much lower edge density than the rule influence diagram with only a slightly larger number of nodes. It also has fewer nodes than conventional and compact rule visualizations because it coarse-grains the specified structures to atomic patterns. We find that the full rule-derived regulatory graph is too dense to permit visual analysis of the functional layer, which only becomes possible when complexity reduction methods are applied. Background removal and node grouping with strict or permissive edge signature provide increasing levels of complexity reduction and enable identification of signaling cascades and network motifs as discussed above. The compressed graphs also approach the size of the contact map while having moderately higher edge densities.

## Discussion

In this work, we have developed new visualizations for the mechanistic, regulatory and functional layers of the rule-based model, namely compact rule-visualization, rule-derived regulatory graph and compressed regulatory graph respectively. We have introduced new data structures to rule-based modeling that enable these visualizations, such as the rule structure graph, the atomic pattern and the edge signature. We have developed a procedure to systematically derive more abstract information such as regulation and function from more concrete layers such as structure and mechanism. The rule-derived regulatory graph and its compressed forms enable automated generation of signaling diagrams for models with hundreds of rules, which was not possible prior to this work. The compression methods can also flexibly account for the nuances of specific biological systems. Graph analysis on the regulatory graph reveals network motifs that are useful for understanding the architecture of the model. Complexity analysis demonstrates that these visualizations are more readable and informative than other automated visualizations such as the contact map [16], the rule influence diagram [17], and the Simmune network viewer [19]. We are hopeful that the rule-derived regulatory graph will play an important role in the transparent documentation of rule-based models and make rule-based modeling accessible to a wider audience.

It is useful to compare the methods developed in this work to other related visualization tools (Fig 14). The Systems Biology Graphical Notation (SBGN) is a set of community standards for visualization of signaling models [20]. Of these, the SBGN Process Description provides a structured representation for reacting entities, so it can be adapted for conventional rule visualization (Fig 14A). A number of other software frameworks also support conventional rule visualization (Simmune [27], Virtual Cell [37,38], BioUML [39]). Compact rule visualization, which we have proposed here, is an alternative to these approaches that has the advantage of conveying the action of a rule explicitly without requiring left-to-right visual comparison. The SBGN Entity Relationship (Fig 14B) [20] and its predecessor the Molecular Interaction Map (Fig 14C) [21] are alternatives to the Extended Contact Map [14], but they also require manual construction by interpreting the rules of the model. In contrast, the rule-derived regulatory graph and its compressed form are automatically generated from the model. The Kappa story (Fig 14D) [16,18] is a visualization that is similar in form to the rule influence diagram [17] and it is generated by simulating possible sequences of rule firings and compressing them internally [18]. While it is useful for showing a specific biochemical trajectory, it requires comparisons of multiple simulation trajectories to build a model diagram. In contrast, the rule-derived regulatory graph of the model is assembled by analyzing individual rules and aggregating them linearly (see Methods).

**Fig 14.**
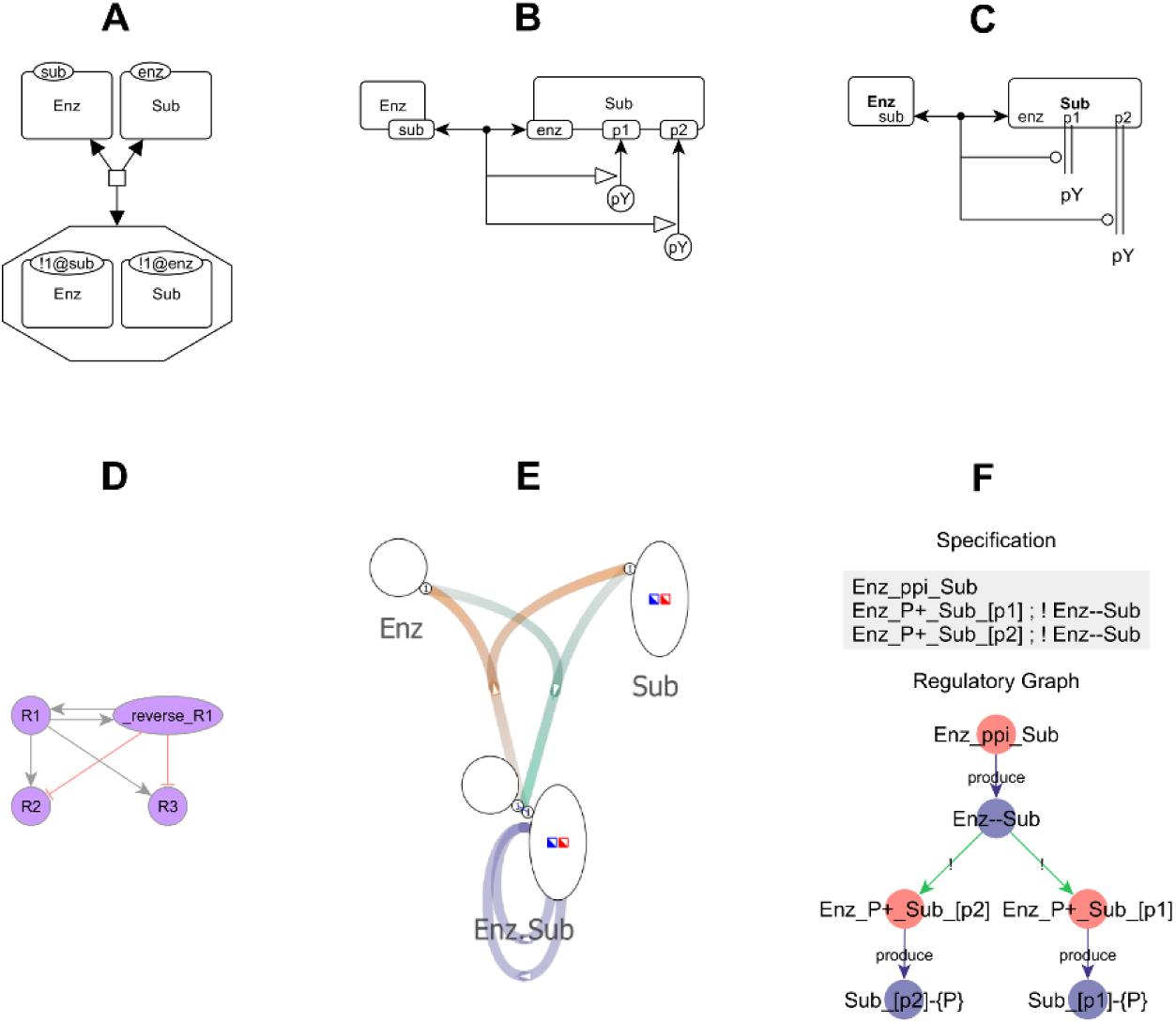
Other visualization tools. **(A)** The SBGN Process Description as applied to conventional rule visualization. **(B)** The SBGN Entity Relationship and **(C)** Molecular Interaction Map are alternatives to the Extended Contact Map. **(D)** The Kappa Story is similar to the rule influence diagram. **(E)** The Simmune Network Viewer produces a compact diagram of rules by merging patterns that differ only in internal states. **(F)** The *rxncon* specification specifies the model itself as a network of regulatory interactions, which is visualized easily as the *rxncon* regulatory graph. See additional notes in Figs S4 and S5.

The Simmune Network Viewer (Fig 14E) [19] visualizes rules as a network in which each node represents a set of molecules connected in a unique manner. The diagram in Fig 14E shows four rules, one modeling enzyme-substrate binding, one modeling dissociation and two modeling phosphorylation of sites on the enzyme-substrate complex. The reactants and products of the phosphorylation rules have the same molecular connectivity (enzyme bound to substrate) but different internal states (unphosphorylated, phosphorylated), so they are represented by the same node in Fig 14E and the paths representing the phosphorylation rules loop back on to that node. This type of compression produces relatively large yet readable diagrams (see data points representing Simmune Network Viewer in Fig 13 and Fig S3), but it obscures signal flow through internal states of molecular components, as we show in Fig. S4. In contrast, the rule-derived regulatory graph does not hide signal flow through any type of structure, including binding sites, bonds and internal states.

Another approach to model construction is to specify regulatory logic using a simple pairwise interaction format and construct mechanisms automatically using a predefined algorithm (Process Interaction Model [40], *rxncon* [22]). In the *rxncon* specification, the model format can be directly translated into a regulatory graph and other related visualizations [22]. However, as we show in Fig S5, the *rxncon* model format is not expressive enough to distinguish between all biological mechanisms that can be modeled as rules and can only reconstruct a restricted set of mechanisms. This motivated the need for a general rule-derived regulatory graph that can be derived from a reaction rule of any size and complexity, such as the rules found in BioNetGen, Kappa and Simmune models.

Although the rule-derived regulatory graph offers a number of advantages over existing methods, there are a number of ways in which it could be improved. Currently, the graph resolves the types of structures that are shared between rules, but it does not explicitly resolve higher order combinations of structures, e.g., there is no separate node that represents the concurrent 3’ and 5’ phosphorylated states on the PIP3 molecule. Similarly, the edge types resolve reactant, product and context relationships, but there could be more complex relationships such as inhibition by consumption of an active state. Also, the current rule grouping algorithm treats each rule as a standalone process, whereas in reality there are modules of processes that work in concert, such as the individual steps of a Michaelis-Menten mechanism. These improvements will be implemented using a combination of graph analysis and user annotation. Finally, BioNetGen supports compartmental states and transport operations [10,41],which are not addressed in the current version of the regulatory graph. We plan to add these features in the future as well as explore alternative graph visualizations, e.g., a matrix representation instead of nodes and edges [36].

Edward R. Tufte, a pioneer of modern data visualization and analytic design, has argued that “cognitive tasks should be turned into design principles”, based on the idea that “universal cognitive tasks” underlie how humans perceive and understand information [42]. A number of such cognitive tasks can be found in diagrams and text-based descriptions that are ubiquitous in the biochemical literature and we recapitulate some of these in our automated methods. First, a biochemical process is usually described as an action one entity performs on another, such as ‘binds’ and ‘phosphorylates’. Graph operation nodes in compact rule visualization (Fig 5B) play a similar role in conveying the action of a rule. Second, the term ‘site’ is commonly used to denote any part of a molecule that behaves distinctly from other parts. A site is also a location at which a kinetic process can occur, e.g. a phospho-motif or binding interface. The definition of atomic patterns in this work (Fig 6A) introduces the notion of actionable sites for rule-based models (similar to elemental states in *rxncon* [22]). Third, in descriptions of pathways, there is selective emphasis on activated states and activation processes at the expense of default states, inactivated states, constitutive deactivation processes and background processes that are not influenced by signal flow. This allows the reader to filter redundant information and focus on what happens after signal is initiated. Background removal on the regulatory graph follows a similar principle (Fig 7B). Fourth, sites and processes are categorized in the literature into families based on homology and functional similarity, and this approach has been effectively used to organize biochemical knowledge [43–46]. The grouping of atomic patterns using user annotation and heuristics (Fig 7C) and the grouping of rules by edge signature (Fig 7D) recapitulates this approach. Finally, generalizing interactions between individuals as an interaction between their families is a common strategy to make broad conclusions about network architecture. Graph compression by collapsing groups (Fig 7E) performs the same function and results in a similarly broad network representation.

The rule-derived regulatory graph is a compact global representation of the rule-based model. It is also a bipartite graph, so it can be provided as input for tools that require simple graphs or bipartite graphs. For example, numeric values from a simulation trace can be mapped to node size or edge thickness on the regulatory graph, resulting in an animated simulation trajectory (e.g. [47]). As *rxncon* developers have shown, it is possible to treat the regulatory graph as a Boolean model and perform stochastic simulations [48]. Model reduction approaches developed for rule-based models have previously used information on regulatory interactions [49] that can now be obtained directly from the regulatory graph. The rule-derived regulatory graph also serves as a rich source of information that could be mined using formal approaches. For example, the identification of model subsystems such as those shown in Figs 10 and 11 could be performed by graph partitioning methods [50]. Network motifs such as those shown in Fig 12 could be identified by cycle detection [51]. Groups of rules and atomic patterns on the regulatory graph could be inferred by structure discovery methods for graph data [52,53]. Thus, the rule-derived regulatory graph paves the way for novel applications of graph analysis, data mining and machine learning to rule-based models.

A natural future direction for signaling models is to integrate rules from different sources and explore the effects of complex input stimuli and crosstalk between pathways [54,55]. Currently, models of signaling from various receptors have as many as hundreds of rules [7–9], and this number is expected to increase by an order of magnitude to cover more molecule types, receptors and signal pathways. Since databases of rules targeting different pathways are being constructed in tandem by different groups (e.g. [7–9,56]), we expect that the complex models of the future will integrate kinetic information from multiple databases. The recently published whole cell model of *Mycoplasma genitalium* [5] makes effective use of databases to organize and visualize kinetic information [57–59] and provides proof-of-concept of a database-oriented approach. We expect that the rule-derived regulatory graph will play a role in building these large models and rule-based databases. A number of exchange formats now support rule-based data structures (SBML-Multi [28], BioPax Level 3 [60]) and a number of modeling frameworks integrate BioNetGen rules with higher-order model composition (Virtual Cell [37,38], PySB [56]). Frameworks that integrate rules can use the rule-derived regulatory graph for visualization and analysis. Sophisticated group structures [61], in addition to the ones implemented so far, will facilitate navigation and visualization of rule-based databases, similar to approaches that have been deployed to analyze other biological data (VisANT [62], ChiBE [63]). Simulations of large models could be optimized by identifying parallelizable submodules [5], and these could be discovered through graph analysis. Thus, in addition to the immediate benefit of visualizing and understanding large models, the rule-derived regulatory graph is expected to be useful in developing the comprehensive cell models of the future.

